# ILC homeostasis phenotyping in various tissues, aging, and sex differences

**DOI:** 10.1101/2021.12.17.473226

**Authors:** Alexis S Mobley, Jesus Bautista Garrido, Pedram Honarpisheh, John d’Aigle, Louise D McCullough, Jaroslaw Aronowski

## Abstract

Innate lymphoid cells are the innate counterpart to CD4+ T cells that are mediated by the same transcription factors and produce similar cytokines. ILCs are being investigated in many different disease states, but the field current lacks foundational information on ILC representation whether it be in tissues, between males and females, or in aging as these are all vital components in disease etiology and severity. Our descriptive study used flow cytometry to characterize ILCs compared to the entire CD45+ (e.g., lymphocyte) and lineage negative (e.g., ILC) compartments to understand their homeostatic balance and plasticity. Moreover, we defined ILC2 expression and subsets based on their cytokine production and created several mathematical models to elucidate the correlation of extra- and intra-cellular ILC2 markers from least to most complex. ILC studies would benefit from more unbiased, holistic experiments including RNA-seq and mass spectroscopy to further define ILCs in steady state before adding more complex pathways like different disease states to enhance translational value and therapeutic targeting of these cells.

## Introduction

Innate lymphoid cells (ILC) are a novel, tissue-resident immune cell that are the innate counterpart to T cells. Historically, we have known about natural killer cells, the CD8+ T cell counterpart, but now we have CD4+ counterparts under the same master transcription factor regulator and cytokine expression: group 1 ILC (ILC1) – Th1, group 2 ILC (ILC2) – Th2, group 3 ILC (ILC3) – Th17/Th22 (1). Though ILCs are rare, they are responsible for pathology in different tissues such as ILC1s in rheumatic arthritis of the joints (2), ILC2s in allergic asthma in the lungs (3), and ILC3s in colitis in the gut (4). However, ILCs are being investigated in ischemic-reperfusion injury (5, 6), cancer (7, 8), and viral infections (9, 10) as therapeutic targets. Current studies have focused on ILCs in their respective tissues, but few have looked at aging or sex differences, and to our knowledge, none have taken a systems biology approach to understand the foundational insights of ILCs.

Every living thing is subjected to aging, whether it be biologic or chronologic (11). Aging is the leading risk factor for most diseases. In aging, there is a chronic low-level inflammation, termed inflammaging, that contributes to disease etiology and there are efforts to reverse inflammaging effects. There are several hallmarks in biologic aging, including immunosenescence, that are critical to investigate in disease models. Very few pre-clinical studies are done in appropriately aged mice, and this phenomenon has started to burden ILC research as well.

Sex differences attribute to disease progression and etiology such as autoimmune disorders being common in females (12, 13). Moreover, studies are starting to tie in the importance of hormonal and gonadal sex on immune cell function (14). But as female animals age, they undergo menopause, which changes their hormones, and therefore, immune cell function. It is imperative that more studies investigate criteria like sex differences and aging for better clinical translations.

Considering the inherent differences in tissue function, aging, and sex differences, we thought it important to investigate ILCs in all three conditions to better understand the foundational differences in their presentation and function.

## Materials and Methods

### Mice

Young (6 months; equivalent to 30-year old human or 12-months; equivalent to 40-50-year old human) and aged (24-26 months; equivalent to 60-75-year old human) male and female C57BL6/J mice were obtained from Jackson laboratory and aged in house to control for dietary and environmental factors. Experimental and control mice were co-housed littermates. All animals were group housed in Tecniplast individually ventilated cage (IVC) racks, fed a commercially available irradiated, balanced mouse diet (no. 5058, LabDiet, St Louis, MO) and provided corncob bedding. Rooms were maintained at 70-73^◦^F and under a 12:12-h light:dark cycle. All animals were maintained specific pathogen free for ectromelia virus, epizootic diarrhea of infant mice, lymphocytic choriomeningitis virus, mouse adenoviruses 1 and 2, mouse hepatitis virus, mouse parvovirus, minute virus of mice, polyomavirus, pneumonia virus of mice, reovirus 3, Theiler meningoencephalomyelitis virus, Sendai virus, cilia-associated respiratory bacillus, Mycoplasma pulmonis, Clostridium piliforme, Encephalitozoon cuniculi, Myocoptes spp., Radfordia spp., Myobia spp., Aspicularis tetraptera, Syphacia muris, and Syphacia obvelata. Animal studies were performed in compliance with the U.S. Department of Health and Human Services Guide for the Care and Use of Laboratory Animals. Animal procedures were performed at an AAALAC accredited facility and were approved by the Animal Welfare Committee at the University of Texas Health Science Center at Houston, TX, USA.

### Flow Cytometry

We are performing previously described methods (15, 16). Briefly, mice were deeply anesthetized by Avertin injection and transcardially perfused with 60mL of sterile, ice-cold phosphate buffered saline (PBS). The organs were removed and processed as described below:

### Brain/Lung

Brain and lung tissue were collected and placed in complete Roswell Park Memorial Institute 1640 (RPMI) medium (Gibco). Brain was mechanically and enzymatically digested in collagenase/dispase (1mg/mL) and DNase (10mg/mL; Roche Diagnostics). Lung tissue was mechanically and enzymatically digested with Collagenase/Hyaluronidase (3000U and 1000U, respectively; Stemcell). Tissues were digested for 45 minutes at 37^◦^C. The cell suspension was filtered through a 70µm filter. Leukocytes were harvested from a 70%/30% and 70%/40% Percoll gradient for brain and lung tissue, respectively. Tissues were washed in PBS, counted with a Countess and placed on ice until ready to be stimulated.

### Bone Marrow

Femurs were removed of all bone and fat and the ends removed with a razor. The marrow was flushed with 5mL RPMI and centrifuged. The marrow was then filtered through a 70µm filter and red blood cells lysed with two rounds of 1X RBC lysis buffer (Tonbo Biosciences). Cells were washed in PBS, counted with a Countess, and placed on ice until ready to be stimulated.

### Spleen

Spleen was pushed through a 70µm filter using the back of a syringe. The filter was washed with RPMI and the cells centrifuged. Splenic cells underwent two rounds of red blood cell lysing using 1X RBC lysis buffer (Tonbo Biosciences). Cells were washed in PBS, counted with a Countess, and placed on ice until ready to be stained.

### Gut

Small and large intestines were rapidly removed and placed in ice-cold PBS. The intestinal tissue was opened longitudinally after removal of fat and connective tissues. Fecal content was removed, and the tissue was cut into approximately 1.0cm pieces after being washed with in ice-cold PBS. Gut tissue was incubated in 5mL 5mM ethylenediaminetetraacetic acid (EDTA) in Hank’s Buffered Salt Solution (HBSS, Invitrogen) for 30 mins at 37^◦^C with 100rpm rotation. The epithelial cell later was removed and filtered through a 70µm cell strainer. The retrieved intestinal pieces were washed in HBSS, cut into smaller pieces, and immersed in 10mL digestion solution containing 5% heat-inactivated fetal bovine serum (FBS), collagenase IV (1.75mg/mL; Roche) and DNase I (0.5mg/mL) at 37^◦^C for 45 minutes at 100rpm rotation. Cells were washed with PBS, counted with a Countess, and placed on ice until ready to be stimulated.

### Stimulation

Cells were plated at 2X10^6^/well on U bottom cell culture plates in RPMI supplemented with 10% heat-inactivated fetal bovine serum (FBS), 100U/mL penicillin, 100µg/mL streptomycin, 10mM HEPES buffer, 1X MEM non-essential amino acids, 1mM sodium pyruvate, 50µm β- mercaptoethanol, and 50µg/mL gentamycin. Cells were stimulated with 20ng/mL of the following cytokines: IL-7 (Tonbo), IL-2/IL-25 (Tonbo/R&D Sciences), or IL-33 (Tonbo Biosciences). Negative control cells were treated with PBS while positive control cells were treated with Cell Stimulation Cocktail (eBioscience). All cells were treated with 1X Protein Transport Inhibitor Cocktail (eBioscience) and incubated overnight at 37^◦^C and 5% CO2.

### Staining

Cells were washed with PBS and stained with 1:1000 Ghost Violet 510 viability dye (Tonbo) for 30 minutes at room temperature (RT) in the dark. Cells were washed with FACS buffer (2% FBS and 1mM EDTA in PBS) then blocked with 1:100 CD16/32 (Thermo) in FACS buffer for 30 minutes at RT. For cell surface analysis, cells were washed with FACS buffer and stained with Brilliant Violet 605-CD45 (Biolegend), FITC-Lineage (Lin), Brilliant Violet 421-IL-23R, PE-Cy7-NKp46, APC-ST2, and AlexaFluor 700-NK1.1.

For stimulation studies, cells were washed with FACS buffer and stained with Brilliant Violet 605-CD45 (Biolegend), PE-ST2 (Biolegend), APC/Fire 750-ICOS (Biolegend), and FITC-Lin cocktail (Biolegend and Tonbo Biosciences; Table 1) for 30 minutes at RT in the dark. Cells were fixed and permeabilized with BD Cytofix/Cytoperm kit. Briefly, cells were fixed with Fixation/Permeabilization buffer for 20 minutes at 4^◦^C. Cells were washed twice with 1X BD Perm/Wash Buffer (PWB) and stained with PE-eFluor610-IL-13 (Invitrogen), Brilliant Violet 421-IL-5 (Biolegend), and APC-IL-4 (Tonbo Bioscience) for 30 minutes at 4^◦^C. Cells were washed two times in BD PWB then resuspended in 250uL of FACS buffer for analysis. All antibodies were titrated and validated before the experiment and appropriate fluorescence minus one (FMOs) and compensation controls made per antibody using cells for viability dyes and OneComp eBeads (Thermo) for all others.

**Table 1.** Antibody details and clones

| Antigen/Stain | Fluorophore | Clone | Company |
| --- | --- | --- | --- |
| Extracellular staining |  |  |  |
| Dead cells | Ghost Dye Violet 510 | N/A | Tonbo Biosciences |
| CD45 | Brilliant Violet 605 | 30-F11 | Biolegend |
| IL-23R | Brilliant Violet 421 | 12B2B64 | Biolegend |
| CD335/NKp46 | PE-Cy7 | 29A1.4 | Biolegend |
| IL-33Rα/ST2 | APC | DIH9 | Biolegend |
| NK1.1 | AlexaFluor 700 | PK136 | Biolegend |
| Lineage Markers |  |  |  |
| CD3 | FITC | 17A2 | Tonbo Biosciences |
| CD4 | FITC | RM4-5 | Tonbo Biosciences |
| CD19 | FITC | 1D3 | Tonbo Biosciences |
| TER-119 | FITC | TER-119 | Tonbo Biosciences |
| CD11b | FITC | M1/70 | Tonbo Biosciences |
| CD45R/B220 | FITC | RA3-6B2 | Tonbo Biosciences |
| CD31 | FITC | 390 | Biolegend |
| F4/80 | FITC | BM8.1 | Tonbo Biosciences |
| Fc R1α | FITC | 1-Mar | Biolegend |
| CD23 | FITC | B3B4 | Biolegend |
| CD49b | FITC | DX5 | Biolegend |
| CD326/Ep-CAM | FITC | G8.8 | Biolegend |
| CD11c | FITC | N418 | Tonbo Biosciences |
| Ly-6G/Gr-1 | FITC | RB6-8C5 | Tonbo Biosciences |
| CD8a | FITC | 53-6.7 | Tonbo Biosciences |
| Intracellular/Stimulation Staining |  |  |  |
| IL-5 | Brilliant Violet 421 | TRFK5 | Biolegend |
| ST2 | PE | DIH9 | Biolegend |
| IL-13 | PE-eFluor610 | eBio13A | Invitrogen |
| IL-4 | APC | 11B11 | Tonbo Biosciences |
| CD278/ICOS | APC/Fire 750 | C398.4 | Biolegend |

A detailed list of antibody/dye targets, fluorophores, clones, and companies.

### Flow Cytometric Analysis

Cells were run on a Beckman Coulter Cytoflex S. FMOs were used to establish gates. 2.5X10^5^ CD45+ cells (or until the well ran dry) were run on medium settings. For the characterization experiments, ILCs were identified as follows: live, singlet, CD45+ Lin-then NK1.1+ NKp46+/-for ILC1, IL23R+ NKp46+/- for ILC3, and NK1.1-IL-23R-ST2+ cells for ILC2s. Group 2 innate lymphoid cells were identified as single, live CD45+/Lin-/ST2+/ICOS+ cells and then gated on IL-4+, IL-5+, and IL-13+ cells. The flow cytometry results were analyzed using FlowJo^TM^ v10.8 Software (BD Life Sciences).

### Statistical Analysis

All analysis was done with GraphPad Prism version 9.0.2 for Mac. Outliers were identified using ROUT (Q = 0.2%) with the average of replicates in each row and then calculated for each column to handle the subcolumns. Outliers were removed and data analyzed. Three-way analysis of variance (ANOVA) with Tukey’s correction for multiple comparisons were performed when appropriate. Repeated measures were implemented for the lymphocyte and ILC compartments. If data were missing for the repeated measures, Prism ran a mixed-effects model instead without assumption of sphericity. Geisser-Greenhouse’s epsilons (GGE) were reported in these cases. If effects were not significant during the three-way ANOVA, data were consolidated and re-analyzed with two-way ANOVA to understand if the previously significant effects were still significant in the data. Multiple linear regression and simple linear regression were performed for the main effects of the data. Non-significant effects were removed to simplify the data model. P values, or adjusted P-values for multiple comparisons, were reported with significance denoted as <0.05.

## Results

### Aging and sex differences mediate ILC phenotypic changes in the spleen

To our knowledge, there have been no ILC comparative studies in mouse models in various tissues in aging and sex differences. We were able to extensively study ILC percentages in the CD45+ lymphocyte compartment (Figure 1A) and ILC compartment (Figure 1B). ILCs are differentially expressed in the spleen [F (2, 36) = 35.83, p<0.0001] where males make more ILCs than females [F (1, 36) = 50.37, p<0.0001]. There are more male young (p<0.0001) and aged (p = 0.0002) ILC1s than females; however, aged males (AM) have more ILC3s than young males (YM; p = 0.0323) and aged females (AF; p = 0.0303). For the young male, the spleen was ILC1-dominant containing more ILC1s than ILC2s (p = 0.0008) and ILC3s (p = 0.0004) and more ILC3s than ILC2s (p = 0.0102).

**Figure 1.**
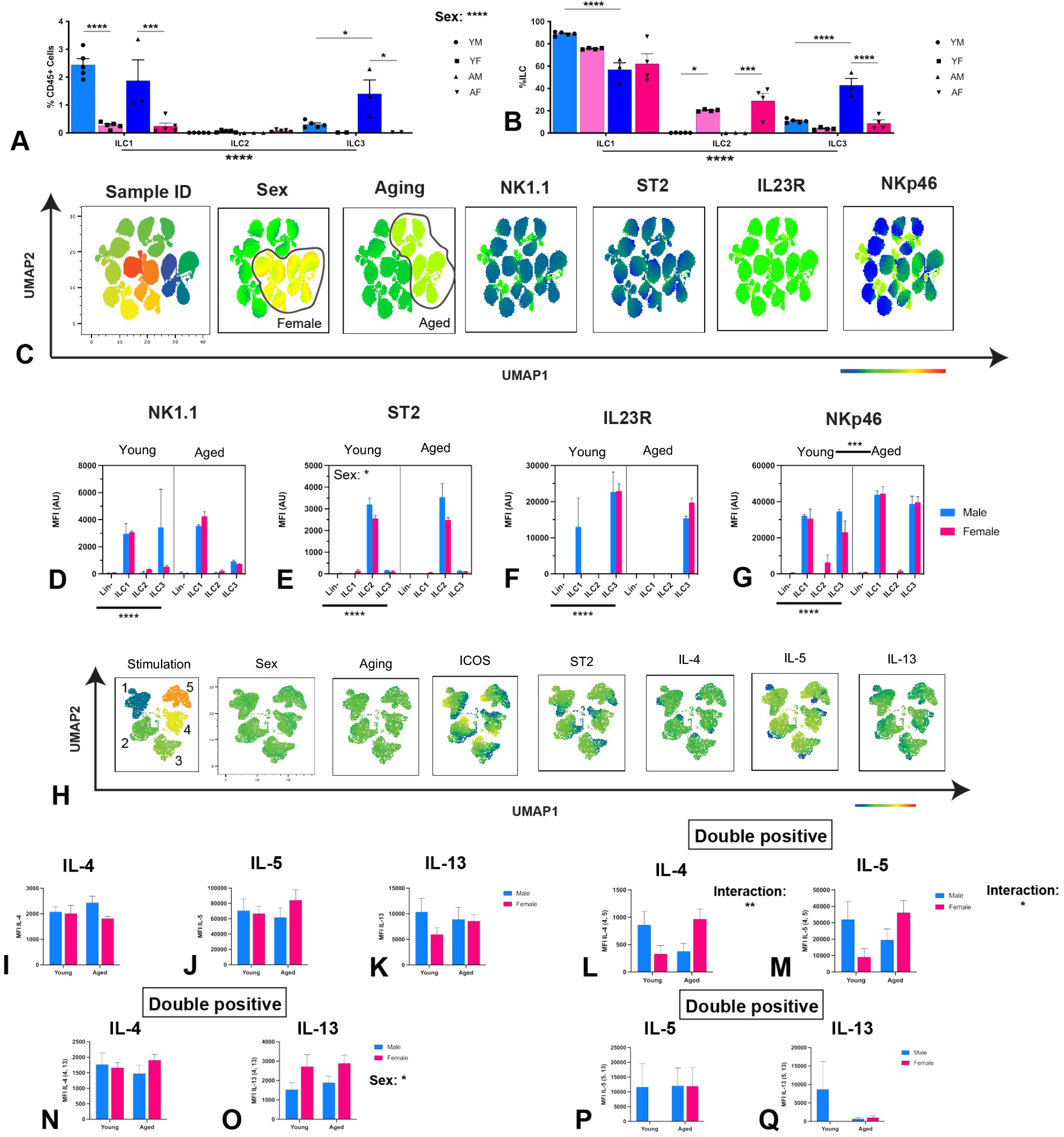
Aging and sex differences influence splenic ILC representation and function. Immune cells were isolated from the spleen of young (12-months) and aged (24-26-months) male and female C57Bl6/J (WT) mice. The percentage that ILCs comprise the CD45+ lymphocytic (A) and ILC (B) compartments are represented and separated by young males (YM), young females (YF), aged males (AM), and aged females (AF). Uniform manifold approximation and projection (UMAP) was utilized for dimensional reduction to understand how the cell surface markers, sex, and aging influenced clustering (C). NK1.1 (D), ST2 (E), IL23R (F), and NKp46 (G) median fluorescence intensity (MFI) were used to assess ILC lineage commitment. Three-way ANOVA with Tukey’s analysis for multiple comparisons was used to determine effect significance and sample differences. If an effect wasn’t significant, data were consolidated to remove that effect and two-way ANOVA were performed (A-B, D-G). Splenic cells of young (6-months) and aged (24-26 months) male and female were isolated, stimulated overnight with PBS (negative control – denoted 1), PMA (positive control – denoted 5) or 20ng/mL IL-7 (denoted 2); IL-2 and IL-25 (denoted 3); or IL-33 (denoted 4), and investigated for type 2 cytokine expression using flow cytometry. ILC2s were isolated and investigated for type 2 cytokine expression using flow cytometry. UMAPs were used to assess similarities between ILC2s and delineate ILC2 cytokine subtypes (H). The cytokine production of single- and double-cytokine positive ILC2s were reported: ILC24 (I), ILC25 (J), ILC213 (K), ILC2_4,5_ (L-M), ILC2_4,13_ (N-O), ILC2_5,13_ (P-Q). Two-way ANOVA was used with Tukey’s analysis for multiple comparisons (I-Q). Adjusted P values were reported. When appropriate, data are represented by mean ± standard error of the mean (SEM); n = 3-5/group. p < 0.05 are significant; *: p < 0.0332; **: p < 0.0021; ***: p < 0.0002; ****: p < 0.0001

Because ILCs comprise such a small subset of lymphocytes and favor plasticity phenotypes, we wanted to investigate how ILCs compare to each other in an ILC compartment (Figure 1B). There was still an ILC difference between the three ILC types [F (2, 36) = 288.5, p<0.0001]; however, there was no significant effect from aging or sex differences. Regardless of major effect changes, there was still a decrease of ILC1 in aging in males (p<0.0001). Young females (YF) and AF had more ILC2s than their male counterparts (p = 0.0152, p = 0.0009, respectively). As seen in the lymphocyte compartment, there were more AM ILC3s than YM (p<0.0001) and AF (p<0.0001). For the male spleen, there is ILC1-dominance compared to ILC2 (p<0.0001) and ILC3 (p<0.0001). This changes in aging where there are more ILC1s than ILC3s (p<0.0001), but more ILC3s than ILC2s (p=0.0004). For females, the spleen is ILC1-dominant compared to ILC2s (p<0.0001) and ILC3s (p<0.0001) and holds true in aging for ILC2s (p = 0.0001) and ILC3s (p<0.0001). Compared to ILC3s, ILC2s increase in aging (p = 0.0263).

Though ILCs can be distinctly categorized, they are uniquely able to express similar receptors and cytokines as their sister cells (17). To visualize this phenomenon, we used uniform manifold approximation and projection (UMAPs) for dimensional reduction to compress the multi-dimensionality of the generated flow cytometry data to understand how extracellular receptors, sex, and aging influence ILC phenotypes (Figure 1C). NK1.1 is a cellular receptor present on natural killer cells and ILC1s, IL-23R for ILC3, and ST2 for ILC2s (18). NKp46 has been identified as a cytotoxicity marker for both ILC1s and ILC3s (18) and was used in this analysis as such. To indicate a pure, committed ILC lineage, it would be anticipated that ILCs are presenting their specific maker exclusively or in significantly higher amounts than their sister cells. We used flow cytometry to assess ILC lineage commitment from cell surface marker median fluorescence intensity (MFI) as influenced by sex and aging in the spleen. Indeed, there was a difference in NK1.1 [F (3, 48) = 10.74, p<0.0001; Figure 1D], ST2 [F (3, 48) =347.9, p<0.0001; Figure 1E], IL23R [F (3, 48) = 35.83, p<0.0001; Figure 1F], and NKp46 [F (3, 48) = 199.2, p<0.0001; Figure 1G] in different ILCs. ST2 also had sex differences [F (1, 48) = 6.381, p = 0.0149] that was still maintained when data were consolidated to remove aging [F (1, 56) = 391.7, p = 0.0131; data not shown]. AM expressed more ST2 than YF (p = 0.0077) and AF (p = 0.0033). Aging increased NKp46 expression in the spleen [F (1, 48) = 13.95, p = 0.0005; Figure 1D], which is supported by NKp46 MFI expression from YF compared to AM (p = 0.0463) and AF (p = 0.0106) increases. There were also trends in age-mediated decreases in IL23R expression, but even with a two-way ANOVA to removed sex differences, this effect never reached significance [F (1, 56) = 3.472, p = 0.0677].

Different tissues require different cytokines depending on their inherent nature. Because ILC2 is the uniquely anti-inflammatory ILC type, we probed classic type 2 cytokines to understand the functional changes that occur in this cell. ILC2s are known to produce IL-4, IL-5, and IL-13 and the combination of these cytokines can yield various outcomes (3, 19–22). We utilized UMAPs to understand how sex, aging, and extracellular surface proteins influence ILC2 type 2 cytokine production. By two-way ANOVA, there was no sex or aging differences in single cytokine expression of IL-4 (Figure 1I), IL-5 (Figure 1J), or IL-13 (Figure 1K) in splenic ILC2s. However, when investigating a specific subset of ILC2s that produce both IL-4 (Figure 1L) and IL-5 (ILC2_4,5_; Figure 1M), there is an interaction where in aging, males decrease [F (1, 84) = 9.097, p = 0.0034] and females increase [F (1, 81) = 6.640, p = 0.0118] their cytokine expression. Interestingly, females produce more IL-13 (Figure 1O) in ILC2_4,13_ than males [F (1, 74) = 6.109, p = 0.0157], but this isn’t seen in IL-4 (Figure 1N) expression. Though splenic ILC2s have potential to produce both IL-5 (Figure 1P) and IL-13 (Figure 1Q), there are no significant differences in their expression values.

### ICOS and ST2 expression predict splenic ILC2 cytokine expression

Current literature has implicated ICOS and ST2 as excitatory expression markers in ILC2s, but to our knowledge, there are no experiments to support this. Therefore, we used simple linear regressions to understand how ICOS (Figures 2A-J) and ST2 (Figures 2K-S) predict cytokine output of all ILC2 subsets. We showed that ICOS is inversely proportional to ST2 (p = 0.0191; Figure 2A), IL-4 (p = 0.0486; Figure 2B), and IL-5 (p = 0.0191; Figure 2C); however, ICOS is proportional to IL-5 in ILC2_5,13_ (p = 0.0355; Figure 2I). There was no correlation for IL-13 (Figure 2D), ILC2_4,5_ (Figure 2E), ILC2_4,13_ (Figure 2G-H), or IL-13 of ILC2_5,13_ (Figure 2J).

**Figure 2.**
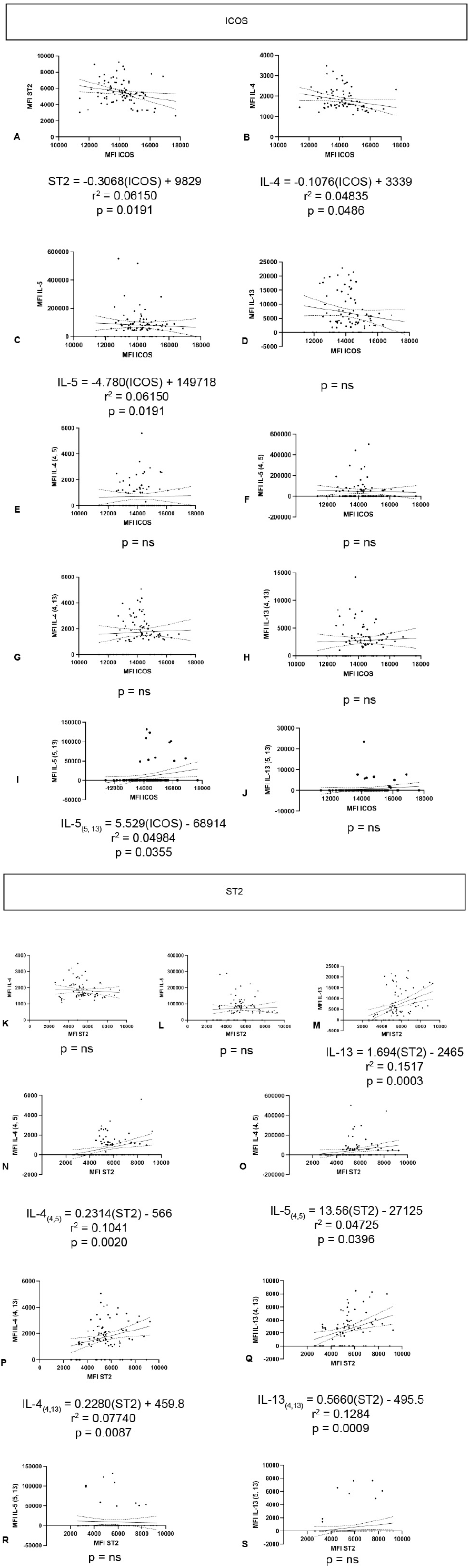
Splenic ILC2 functional output is predicted by cell surface markers. Splenic cells of young (6-months) and aged (24-26 months) male and female were isolated, stimulated overnight with PBS, IL-7, IL-2 and IL-25, IL-33 or, PMA, and investigated for type 2 cytokine expression using flow cytometry. Single linear regressions were graphed with 95% confidence intervals for ICOS against ST2 (A), IL-4 (B), IL-5 (C), IL-13 (D), ILC2_4,5_ (E-F), ILC2_4,13_ (G-H), and ILC2_5,13_ (I-J) and ST2 against IL-4 (K), IL-5 (L), IL-13 (M), ILC2_4,5_ (N-O), ILC2_4,13_ (P-Q), and ILC2_5,13_ (R-S). Only significant models were reported with r^2^ and p-values.

Interestingly, ST2 predicted the metrics that ICOS did not. ST2 was positively correlated to IL-13 (p = 0.0003), IL-4 (p = 0.0020) and IL-5 (p = 0.0396) of ILC2_4,5_, and IL-4 (p = 0.0087) and IL-13 (p = 0.0009) of ILC2_4,13_. There was no correlation for IL-4 (Figure 2K), IL-5 (Figure 2L), or IL-5 (Figure 2R) and IL-13 (Figure 2S) of ILC2_5,13_. In conclusion, splenic ILCs favor ILC1, thought there are some plastic ILC1/3. Cytotoxicity increases in aging as indicated by NKp46 MFI expression values. Moreover, all ILC2 cytokine subsets, expect for ILC24, 5, 13, are represented in the spleen and ICOS and ST2 predict cytokine expression in ILC2s.

### The bone marrow favors ILC1, which may be mediated by hematopoiesis

As in the spleen, we investigated ILC representation in the lymphocyte (Figure 3A) and ILC (Figure 3B) compartments in aging and sex differences. There is ILC differential expression in the bone marrow [F (2, 39) = 23.28, p<0.0001], where males make more ILCs than females [F (1, 39) = 40.31, p<0.0001]. Particularly for ILC1, YM have more ILC1s than YF (p<0.0001) and AM have more ILC1s than YF (p<0.0001). There is still ILC differences in the bone marrow [F (2, 36) = 129.0, p<0.0001]; however, there is no aging or sex differences, and neither can be reconciled with a two-way ANOVA. As seen in the lymphocyte compartment, YM have more ILC1s than YF (p<0.0001) and AM have more ILC1s than AF (p<0.0001), but YF have more ILC2s than YM (p<0.0001) and AF have more ILC2s than AM (p<0.0001). YM bone marrow has more ILC1s than ILC2s (p = 0.0072) and ILC3s (p = 0.0035) but more ILC2s than ILC3s (p = 0.0035). Once the mouse ages though, there are more ILC1s than ILC3s (p = 0.0418), but ILC2 and ILC3 have similar amounts. Females are ILC2-dominant in the bone marrow compared to ILC1 in young (p<0.0001) and aged (p<0.0001) mice and ILC3s in young (p = 0.0216) and aged (p = 0.0306) mice. There are more ILC2s in the female bone marrow compared to ILC3s in young (p<0.0001) and aged (p<0.0001) mice.

**Figure 3.**
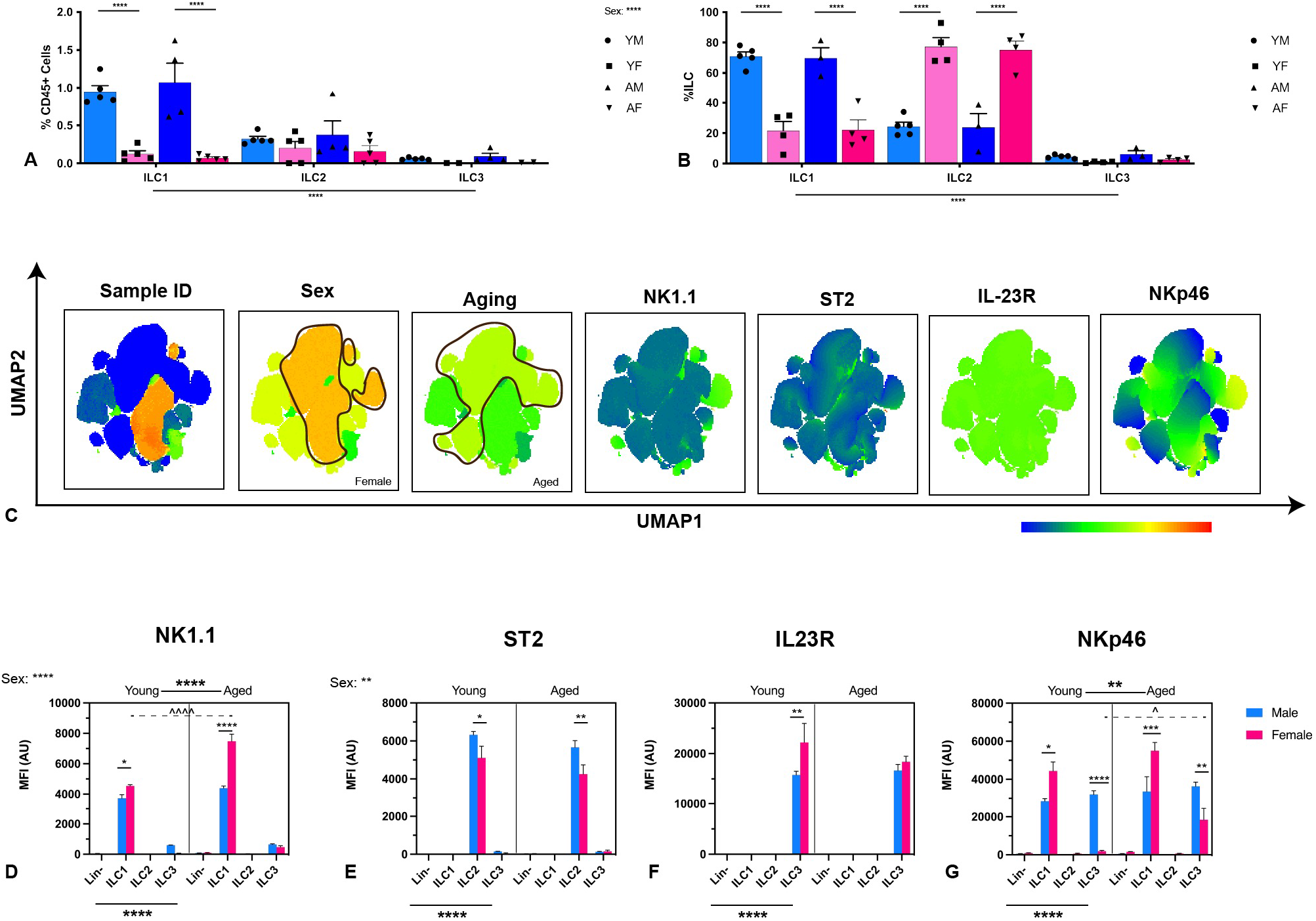
Bone marrow-resident ILCs favor ILC1 in males and ILC2 in females. Immune cells were isolated from the bone marrow of young (12-months) and aged (24-26-months) male and female C57Bl6/J mice. The percentage that ILCs comprise the CD45+ lymphocytic (A) and ILC (B) compartments are represented and separated by young males (YM), young females (YF), aged males (AM), and aged females (AF). Uniform manifold approximation and projection (UMAP) was utilized for dimensional reduction to understand how the cell surface markers, sex, and aging influenced clustering (C). NK1.1 (D), ST2 (E), IL23R (F), and NKp46 (G) median fluorescence intensity (MFI) were used to assess ILC lineage commitment. Three-way ANOVA with Tukey’s analysis for multiple comparisons was used to determine effect significance and sample differences. If an effect wasn’t significant, data were consolidated to remove that effect and two-way ANOVA were performed (A-B, D-G). When appropriate, data are represented by mean ± standard error of the mean (SEM); n = 3-5/group. p < 0.05 are significant; Main effects: *: p < 0.0332; **: p < 0.0021; ***: p < 0.0002; ****: p < 0.0001; multiple comparisons: ^: p < 0.0332; ^^: p < 0.0021.

We utilized UMAPs to visualize dimensionality reduction in the bone marrow (Figure 3C) and quantified NK1.1 (Figure 3D), ST2 (Figure 3E), IL23R (Figure 3F), and NKp46 (Figure 3G). NK1.1 was predominantly expressed in ILC1s [F (3, 60) = 1110, p<0.0001], where females had more NK1.1 expression [F (1, 60) = 51.66, p<0.0001] and increased in aging [F (1, 60) = 29.28, p<0.0001]. Indeed, YF expressed more NK1.1 than YM (p = 0.0142) and similarly seen in aging where AF expressed more NK1.1 than AM (p<0.0001). As supported by the aging effect, AF expressed more NK1.1 than YF (p<0.0001). ST2 also had ILC2-predominant expression [F (3, 60) = 548.9, p<0.0001], where males express more ST2 than females [F (1, 60) = 8.732, p = 0.0045], which is maintained when removing the aging component from the analysis [F (1, 68) = 8.792, p = 0.0042]. This is supported by YM producing more ST2 than YF (p = 0.0195), and AM producing more ST2 than AF (p = 0.0064). IL23R is also expressed exclusively on ILC3s [F (3, 60) = 281.3, p<0.0001]. There were trends towards sex differences in IL23R expression (p = 0.0663), but even with a two-way ANOVA, there was no significance (p = 0597). Regardless, YF expressed more IL23R than YM (p = 0.0058), but this effect was lost in aging. NKp46 was expressed on both ILC1s and ILC3s [F (3, 60) = 167.5, p<0.0001] and aging increases its expression [F (1, 60) = 10.12, p = 0.0023], which is maintained even when removing sex as a factor [F (1, 68) = 4.122, p = 0.0463]. Interestingly, females express more NKp46 on their ILC1s than males whether they are young (p = 0.0152) or aged (p = 0.0006). This effect is reversed in ILC3s though where males produce more NKp46 than their young (p<0.0001) or aged (p = 0.0092) female counterparts. However, there is an aging increase in NKp46 in females (p = 0.0119). In summary, the bone marrow has sex-specific ILC representation and cell surface markers. Males are ILC1-dominant, while females are ILC2-dominant. Moreover, though cell surface expression is ILC-appropriate, there are sex differences in cell surface markers, indicating functional differences in bone marrow resident ILCs.

### Lung-resident ILCs favor ILC2s and contains all ILC2-cytokine-specific subsets

We again investigated the lymphocyte (Figure 4A) and ILC (Figure 4B) compartments in the lung of young and aged, male and female mice. Within the lymphocyte compartment, there were differences in ILC variation [F (2, 36) = 5.990, p = 0.0057], but aging and sex do not influence their representation. This is also seen within the ILC compartment [F (2, 36) = 89.15, p<0.0001]. There were more ILC2s than ILC1s in young (p = 0.0234) and aged (p = 0.0151) male lungs and young (p = 0.0151) and aged (p = 0.0386) male brains, but similar ILC representation for all others. As anticipated, the male lung is ILC2-dominant with more ILC2s than ILC1s in young (p<0.0001) and aged (p<0.0001) lung and ILC3s in young (p = 0.0032) and aged (p<0.0001) lung. ILC3s have higher percentages in the lung than ILC1s (p = 0.0022), but an increase in ILC1s with aging causes the percentages to be the same. Furthermore, ILC2s are the predominant cell type in the female lung compared to ILC1s in young (p <0.0001) and aged (p<0.0001) mice and ILC3s in young (p<0.0001) and aged (p<0.0001) mice, with more ILC1s than ILC3s in young (p = 0.0015) and aged (p = 0.0174) mice. Our UMAPs grouped the samples according to sex and aging differences (Figure 4C). We quantified the cell surface markers MFI and as anticipated, had ILC-specific representation for NK1.1 [F (3, 48) = 23.29, p<0.0001; Figure 4D], ST2 [F (3, 48) = 66.14, p<0.0001; Figure 4E], IL23R [F (3, 48) = 35.83, p<0.0001; Figure 4F], and NKp46 [F (3, 48) = 199.2, p<0.0001; Figure 4G]. There is an aging-mediated increase in IL23R in ILC3s [F (1, 48) = 5.666, p = 0.0213], which is still maintained when removing sex-differences (p = 0.0279). There is evidence of ILC1/2 cells in the lung based on the NK1.1 MFI increase in ILC2s in AF.

**Figure 4.**
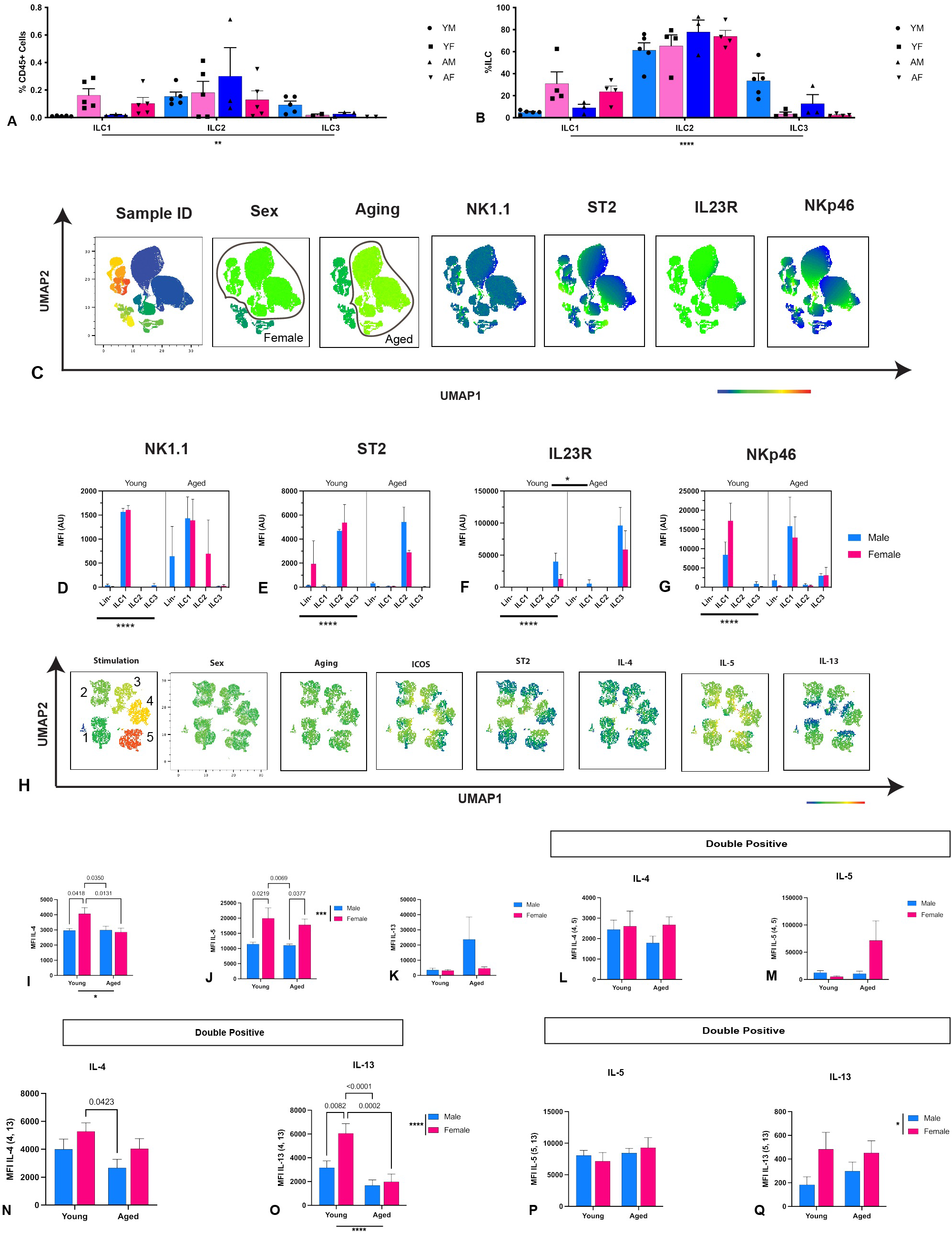
Pulmonary ILCs are lineage-committed but display the most ILC2 cytokine subtypes. Immune cells were isolated from lung from young (12-months) and aged (22-24 month) male and female C57Bl6/J (WT) mice. The percentage that ILCs comprise the CD45+ lymphocytic (A) and ILC (B) compartments are represented and separated by young males (YM), young females (YF), aged males (AM), and aged females (AF). Uniform manifold approximation and projection (UMAP) was utilized for dimensional reduction to understand how the cell surface markers, sex, and aging influenced clustering (C). NK1.1 (D), ST2 (E), IL23R (F), and NKp46 (G) median fluorescence intensity (MFI) were used to assess ILC lineage commitment. Three-way ANOVA with Tukey’s analysis for multiple comparisons was used to determine effect significance and sample differences. If an effect wasn’t significant, data were consolidated to remove that effect and two-way ANOVA were performed (A-B, D-G). Pulmonary cells of young (6-months) and aged (24-26 months) male and female were isolated, stimulated overnight with PBS (negative control – denoted 1), PMA (positive control – denoted 5) or 20ng/mL IL-7 (denoted 2); IL-2 and IL-25 (denoted 3); or IL-33 (denoted 4), and investigated for type 2 cytokine expression using flow cytometry. ILC2s were isolated and investigated for type 2 cytokine expression using flow cytometry. UMAPs were used to assess similarities between ILC2s and delineate ILC2 cytokine subtypes (H). The cytokine production of single- and double-cytokine positive ILC2s were reported: ILC24 (I), ILC25 (J), ILC213 (K), ILC2_4,5_ (L-M), ILC2_4,13_ (N-O), ILC2_5,13_ (P-Q). Two-way ANOVA was used with Tukey’s analysis for multiple comparisons (I-Q). Adjusted P values were reported. When appropriate, data are represented by mean ± standard error of the mean (SEM); n = 3-5/group. p < 0.05 are significant; *: p < 0.0332; **: p < 0.0021; ***: p < 0.0002; ****: p < 0.0001

We utilized UMAPs to visualize the cell surface markers and cytokine potential in pulmonary ILC2s (Figure 4H). Though the samples could cluster by stimulation, there was no distinct clustering based on sex or aging. Aging decreases IL-4 production in pulmonary ILC2s [F (1, 86) = 4.617, p = 0.0345; Figure 4I]. Moreover, YF produce more IL-4 than YM (p = 0.0418), AM (p = 0.0350) and AF (p = 0.0131). When investigating IL-5 (Figure 4J), females produced more IL-5 than their male counterparts [F (1, 79) = 16.10, p = 0.0001], where YF produce more than YM (p = 0.0219) and AM (p = 0.0069), and AF produce more than AM (p = 0.0377). There were no statistical differences in IL-13 or IL-4 and IL-5 in ILC2_4,5_. Surprisingly, there are both aging [F (1, 70) = 20.98, p<0.0001] and sex differences [F (1, 70) = 6.903), p = 0.0106] in IL-13 expression of ILC2_4,13_ (Figure O) but not IL-4 (Figure N). Nevertheless, YF produce more IL-4 than AM (p = 0.0423) for IL-4 and YM (p = 0.0082), AM (p<0.0001), and AF (p = 0.0002) for IL-13 in ILC2_4,13_.

Lastly, females express more IL-13 than males in ILC2_5,13_ [F (1, 74) = 4.898, p = 0.0300; Figure 4Q], but there were no differences in IL-5 expression (Figure 4P).

### Pulmonary ILC2 function is predicted by ICOS and ST2 expression

As in the other tissues, we did simple linear regression to determine how ICOS (Figures 5A-J) and ST2 (Figures 5K-S) correlate to type 2 cytokine expression in ILC2s. IL-4 (Figure 5B), IL-13 (Figure 5D), ILC2_4,5_ (Figure 5E-F), and ILC2_4,13_ (Figure 5G-H) had no correlation to ICOS. We did denote a positive correlation between ICOS and ST2 (p = 0.0005; Figure 5A), IL-5 (p = 0.0009; Figure 5C), and IL-5 of ILC2_5,13_ (p = 0.0204; Figure 5I). Although there was a negative correlation between ICOS and IL-13 for ILC2_5,13_ (p = 0.0162; Figure 5J). ST2 does not correlate with IL-4 (Figure 5K), IL-5 (Figure 5L), IL-13 (Figure 5M), IL-5 of ILC2_4,5_ (Figure 5O), or IL-5 (Figure 5R) and IL-13 (Figure 5S) of ILC2_5,13_. However, ST2 does correlate with IL-4 of ILC2_4,5_ (p = 0.0021; Figure 5N), IL-4 of ILC2_4,13_ (p = 0.0016; Figure 5P), and IL-13 of ILC2_4,13_ (p = 0.0492; Figure 5Q). Based on our analysis, there is no reliable predictor of single cytokine expressing ILC2s, but ICOS and ST2 can be used for pulmonary ILC2 cytokine subsets.

**Figure 5.**
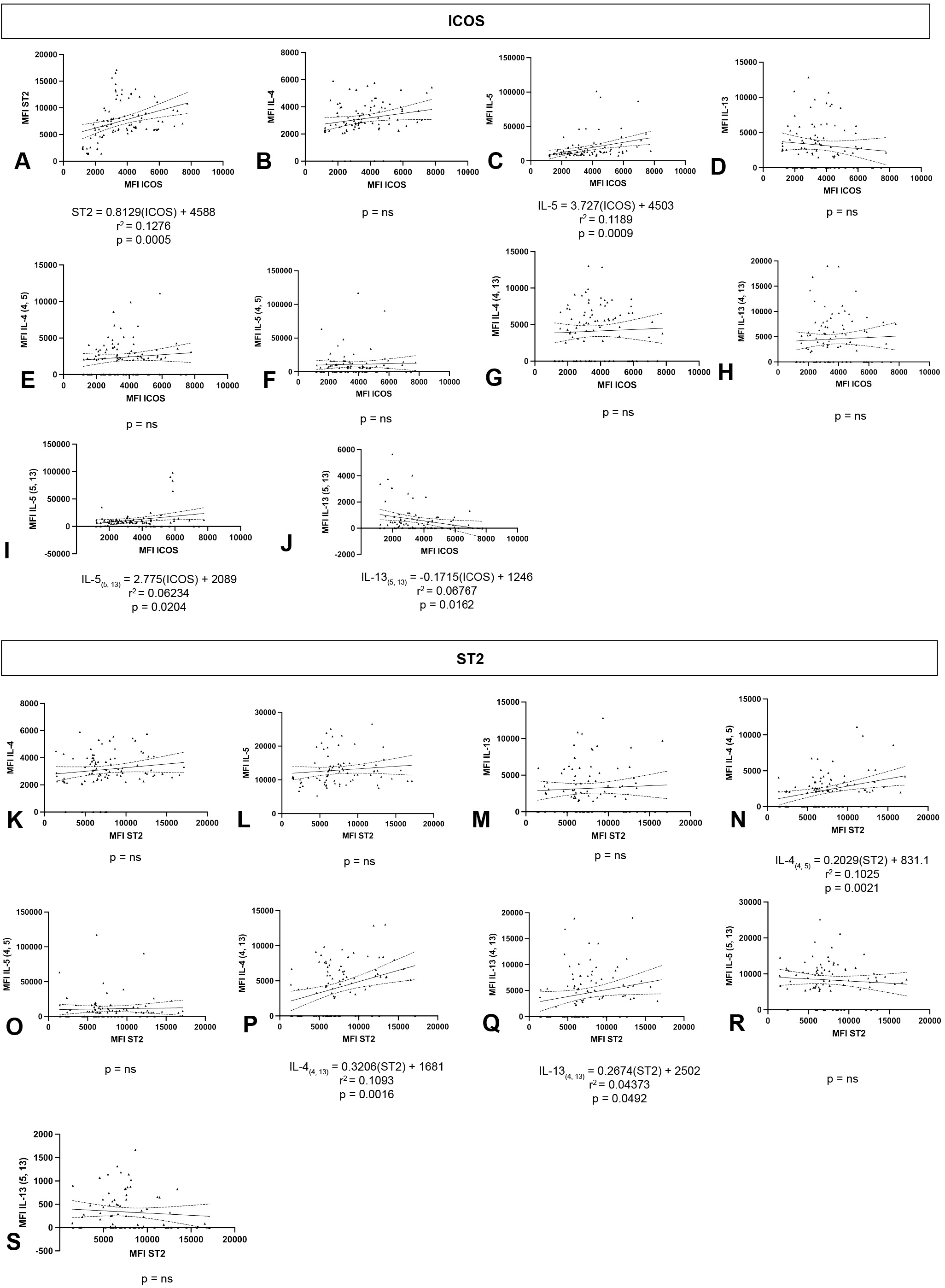
ICOS and ST2 are not great correlates to pulmonary ILC2 function. Pulmonary cells of young (6-months) and aged (24-26 months) male and female were isolated, stimulated overnight with PBS, IL-7, IL-2 and IL-25, IL-33, or PMA, and investigated for type 2 cytokine expression using flow cytometry. Single linear regressions were graphed with 95% confidence intervals for ICOS against ST2 (A), IL-4 (B), IL-5 (C), IL-13 (D), ILC2_4,5_ (E-F), ILC2_4,13_ (G-H), and ILC2_5,13_ (I-J) and ST2 against IL-4 (K), IL-5 (L), IL-13 (M), ILC2_4,5_ (N-O), ILC2_4,13_ (P-Q), and ILC2_5,13_ (R-S). Only significant models were reported with r^2^ and p-values.

### Female gut ILC2s are strongly anti-inflammatory over male ILC2s

For this study, we only investigated ILC2 function in aging and sex differences. Using UMAPs, we clustered the stimulation data and were able to get preferential clustering based on stimulation, sex, and aging (Figure 6A). IL-4 expression (Figure 6B) had both aging [F (1, 83) = 24.96, p<0.0001] and sex differences [F (1, 83) = 11.70, p = 0.0010], where AF produced the most IL-4 over YM (p<0.0001), YF (p = 0.0001), and AM (p = 0.0039). Though IL-5 (Figure 6C) and IL-13 (Figure 6D) did not have aging differences, females continued to produce the most IL-5 [F (1, 75) = 20.70, p<0.0001] and IL-13 [F (1, 84) = 14.60, p = 0.0003]. Indeed, YF produced more IL-5 than YM (p = 0.0429) and AM (p = 0.0213), and AF produced more IL-5 than YM (p = 0.0041) and AM (p = 0.0013). Moreover, compared to AM, YF (p = 0.0076) and AF (p = 0.0028) expressed more IL-13. We investigated ILC2_4,13_ subset (Figures 6E-F), where males failed to produce detectable amounts of either IL-4 [F (1, 69) = 80.06; p<0.0001] or IL-13 [F (1, 73) = 32.69, p<0.0001]. There was an aging-mediated increase in IL-4 in ILC2_4,13_ in females [F (1, 69) = 17.41, p<0.0001], where AF produce the most IL-4. Moreover, IL-13 expressed in YF ILC2_4,13_ was more than YM (p = 0.0148) and AM (p = 0.0196), and AF was more than YM (p<0.0001) and AM (p<0.0001). Lastly, we investigated ILC2_5,13_ where there were no differences in IL-5 expression (Figure 6G); however, there was a sex difference [F (1, 74) = 4.898, p = 0.0300] in IL-13 expression (Figure 6H), where YF produced more IL-13 than YM (p = 0.0430) and AF (p = 0.0231). Based on our findings, female gut ILC2s are particularly active over male ILC2s. There are aging mediated increases in ILC2 function, but it does not have as much of an effect as sex differences.

**Figure 6.**
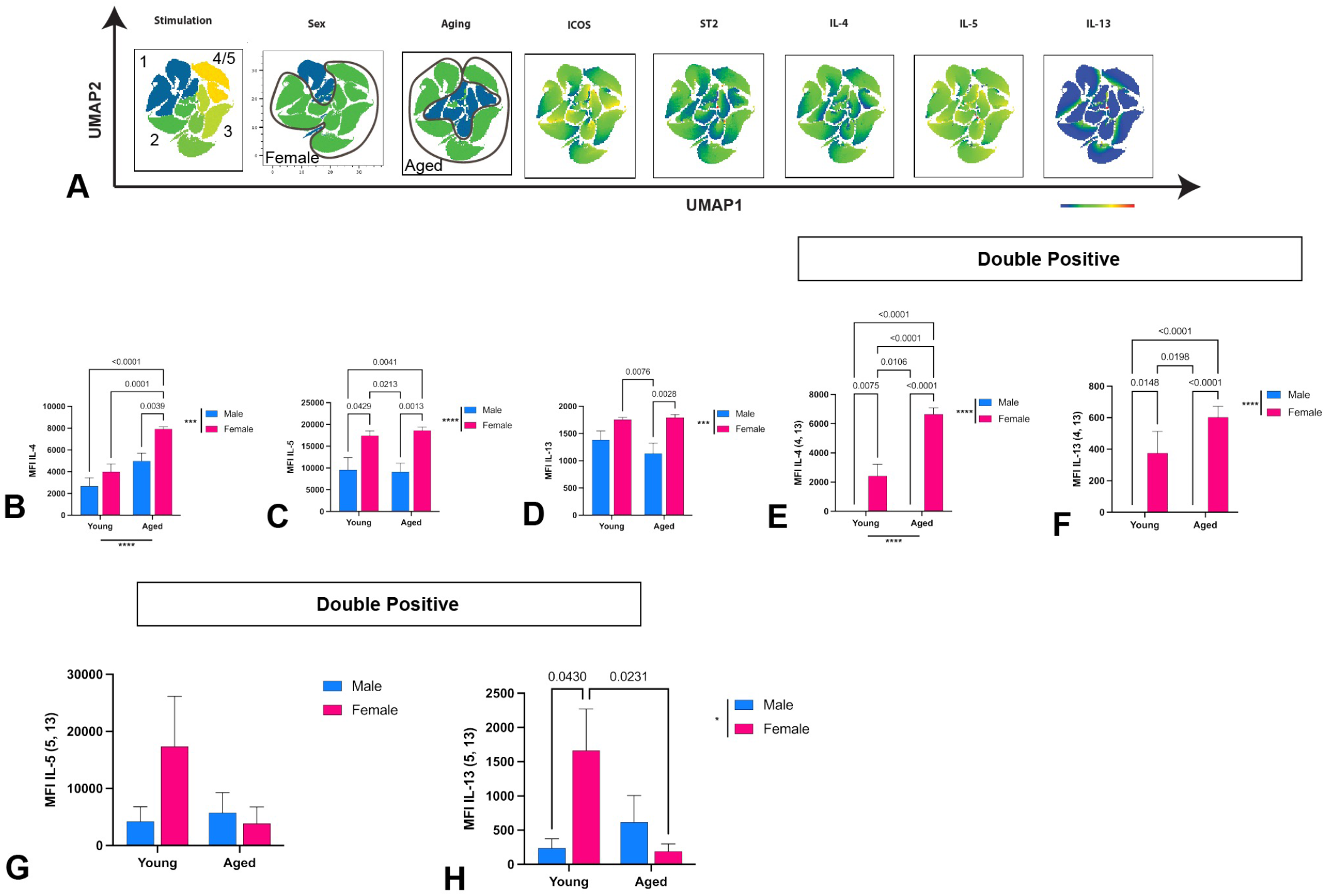
Female gut ILC2s are uniquely anti-inflammatory compared to males. Gut cells of young (6-months) and aged (24-26 months) male and female were isolated, stimulated overnight with PBS (negative control – denoted 1), PMA (positive control – denoted 5) or 20ng/mL IL-7 (denoted 2); IL-2 and IL-25 (denoted 3); or IL-33 (denoted 4), and investigated for type 2 cytokine expression using flow cytometry. CD45+/Lin-/ST2+/ICOS+ ILC2s were isolated and investigated for type 2 cytokine expression using flow cytometry. UMAPs were used to assess similarities between ILC2s and delineate ILC2 cytokine subtypes (A). The cytokine production of single- and double-cytokine positive ILC2s were reported: ILC24 (B), ILC25 (C), ILC213 (D), ILC2_4,13_ (E-F), and ILC2_5,13_ (G-H). Two-way ANOVA was used with Tukey’s analysis for multiple comparisons (B-H). Adjusted P values were reported. When appropriate, data are represented by mean ± standard error of the mean (SEM); n = 3-5/group. p < 0.05 are significant; *: p < 0.0332; **: p < 0.0021; ***: p < 0.0002; ****: p < 0.0001

### Gut activation is negatively correlated with cell surface markers

When probing activation markers for gut ILC2s, no matter if it be ICOS (Figures 7A-H) or ST2 (Figures 7I-O), all statistically significant markers were negatively correlated with activation. IL-5 (p = 0.0121; Figure 7C), IL-13 (p = 0.0380; Figure 7D), and IL-13 of IL2_4,13_ (p = 0.0164; Figure 7F) were correlated with ICOS expression. We also showed that ST2 correlated with IL-4 (p = 0.0271; Figure 7I) and IL-13 of IL2_4,13_ (p = 0.0164; Figure 7M). At the very least, we have reliable activation markers for the single positive ILC2 subsets in this tissue.

**Figure 7.**
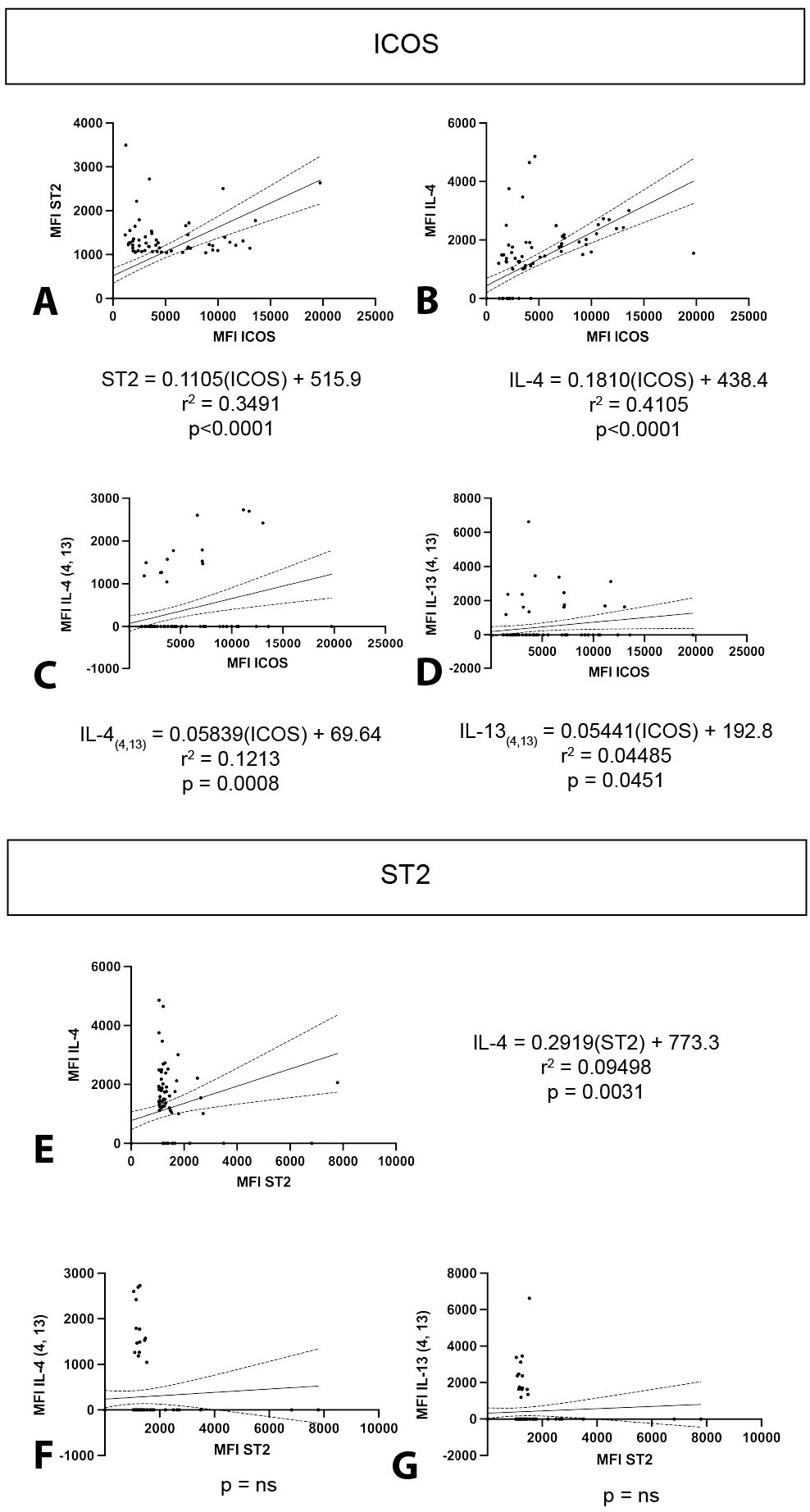
ICOS and ST2 negatively correlate with gut ILC2 activation. Gut cells of young (6-months) and aged (24-26 months) male and female were isolated, stimulated overnight with PBS, IL-7, IL-2 and IL-25, IL-33, or PMA, and investigated for type 2 cytokine expression using flow cytometry. Single linear regressions were graphed with 95% confidence intervals for ICOS against ST2 (A), IL-4 (B), IL-5 (C), IL-13 (D), ILC2_4,13_ (E-F), and ILC2_5,13_ (G-H) and ST2 against IL-4 (I), IL-5 (J), IL-13 (K), ILC2_4,13_ (L-M), and ILC2_5,13_ (N-O). Only significant models were reported with r^2^ and p-values.

### Neural ILCs are uniquely homogenous compared to other tissues

When analyzing neural ILC2s, there was no difference in ILC representation in either the lymphocytic (Figure 8A) or ILC (Figure 8B) compartments. There are aging [F (1, 36 = 12.06, p = 0.0014] and sex-mediated differences [F (1, 36) = 14.99, p = 0.0004] in ILC representation in CD45+ cells, though. AM have more ILC1s than YM (p = 0.0001) and AF (p = 0.0001). There are no major influencers in the ILC compartment, but there are differences between different populations. For example, AM have more ILC1s than AF (p = 0.0085); AF have more ILC2s than YF (p = 0.0117) and AM (p<0.0001); and YM have more ILC3s than YF (p = 0.0361). There were more ILC2s than ILC1s in young (p = 0.0151) and aged (p = 0.0386) male brains, but similar ILC representation for all other ILC types. In females, ILC2s are dominant compared to ILC1s (p<0.0001) and ILC3s (p<0.0001). Taken together, the brain is an ILC2-dominant tissue.

**Figure 8.**
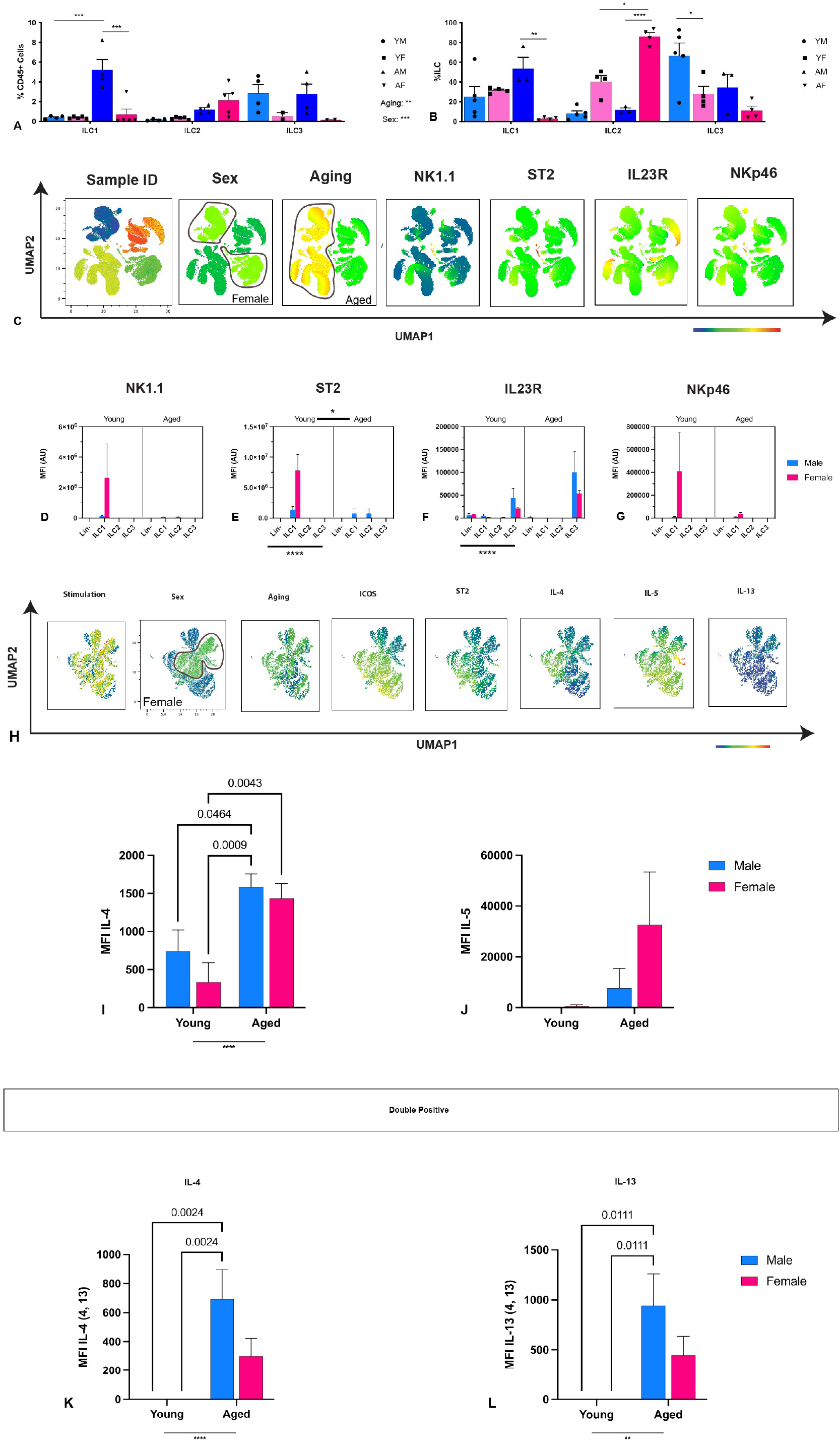
Neural ILCs are homogenous in representation and specific ILC2 cytokine subsets. Immune cells were isolated from the brain of young (12-months) and aged (22-24 month) male and female C57Bl6/J (WT) mice. The percentage that ILCs comprise the CD45+ lymphocytic (A) and ILC (B) compartments are represented and separated by young males (YM), young females (YF), aged males (AM), and aged females (AF). Uniform manifold approximation and projection (UMAP) was utilized for dimensional reduction to understand how the cell surface markers, sex, and aging influenced clustering (C). NK1.1 (D), ST2 (E), IL23R (F), and NKp46 (G) median fluorescence intensity (MFI) were used to assess ILC lineage commitment. Three-way ANOVA with Tukey’s analysis for multiple comparisons was used to determine effect significance and sample differences. If an effect wasn’t significant, data were consolidated to remove that effect and two-way ANOVA were performed (A-B, D-G). Cells of young (6-months) and aged (24-26 months) male and female were isolated from the brain, stimulated overnight with PBS, IL-7, IL-2 and IL-25, IL-33, or PMA, and investigated for type 2 cytokine expression using flow cytometry. ILC2s were isolated and investigated for type 2 cytokine expression using flow cytometry. UMAPs were used to assess similarities between ILC2s and delineate ILC2 cytokine subtypes (H). The cytokine production of single- and double-cytokine positive ILC2s were reported: ILC24 (I), ILC25 (J), and ILC2_4,13_ (K-L). Two-way ANOVA was used with Tukey’s analysis for multiple comparisons (I-L). Adjusted P values were reported. When appropriate, data are represented by mean ± standard error of the mean (SEM); n = 3-5/group. p < 0.05 are significant; *: p < 0.0332; **: p < 0.0021; ***: p < 0.0002; ****: p < 0.0001

We utilized UMAPs to visualize clustering of cell surface markers to investigate ILC plasticity (Figure 8C), where the brain clustered on sex and aging. There was only ILC differentiation for ST2 [F (3, 56) = 9.515, p<0.0001; Figure 6E] and IL23R [F (3, 56) = 19.81, p<0.0001; Figure 8F] when quantified. Neither NK1.1 nor NKp46 had any other effects from sex or aging. ST2 had aging mediated declines [F (1, 56) = 5.906, p = 0.0183]; however, this effect is trending once removing sex as an effect (p = 0.0500). Moreover, ST2 has the strongest representation on ILC1 than ILC2s in young mice, but evens out between ILC1s and ILC2s in aging, suggesting an ILC1/2 hybrid cell.

We also utilized UMAPs to visualize ILC2 clustering (Figure 8H), where ILC2s only clustered on sex, but not stimulation or aging. Brain-resident ILC2s did not produce IL-13 but produced IL-4 and IL-5. ILC24 have an age-related increase in IL-4 expression [F (1, 86) = 18.0, p<0.0001; Figure 8I]. Indeed, AM produce more IL-4 than YM (p = 0.0464) and YF (p = 0.0009), and AF produce more IL-4 than their young counterparts (p = 0.0043). There were no statistical differences in IL-5 expression and no influence from sex or aging (Figure 8J). Interestingly, neural ILC2s could not produce IL-13 alone but only in tandem with IL-4. For both IL-4 (Figure 8K) and IL-13 (Figure 8L), there were aging-mediated increases in ILC2_4,13_ cytokine expression [F (1, 86) = 13.73, p = 0.0004; F (1, 86) = 10.89, p = 0.0014, respectively]. In both cases, AM produced more IL-4 or IL-13 than YM (p = 0.0024, p = 0.0111) and YF (p = 0.0024, p = 0.0111).

### Brain-resident ILC2s have cell surface activation markers

As with the other tissues, we aimed to correlate ICOS (Figure 9A-D) and ST2 (Figure 9E-G). ICOS positively correlated with ST2 (p<0.0001), IL-4 (p<0.0001), and IL-4 (p = 0.0008) and IL-13 (p = 0.0451) of ILC2_4,13_. ST2 was only correlated with IL-4 expression (p = 0.0031). Both ICOS and ST2 can be used as reliable activation markers for brain-resident ILC2s.

**Figure 9.**
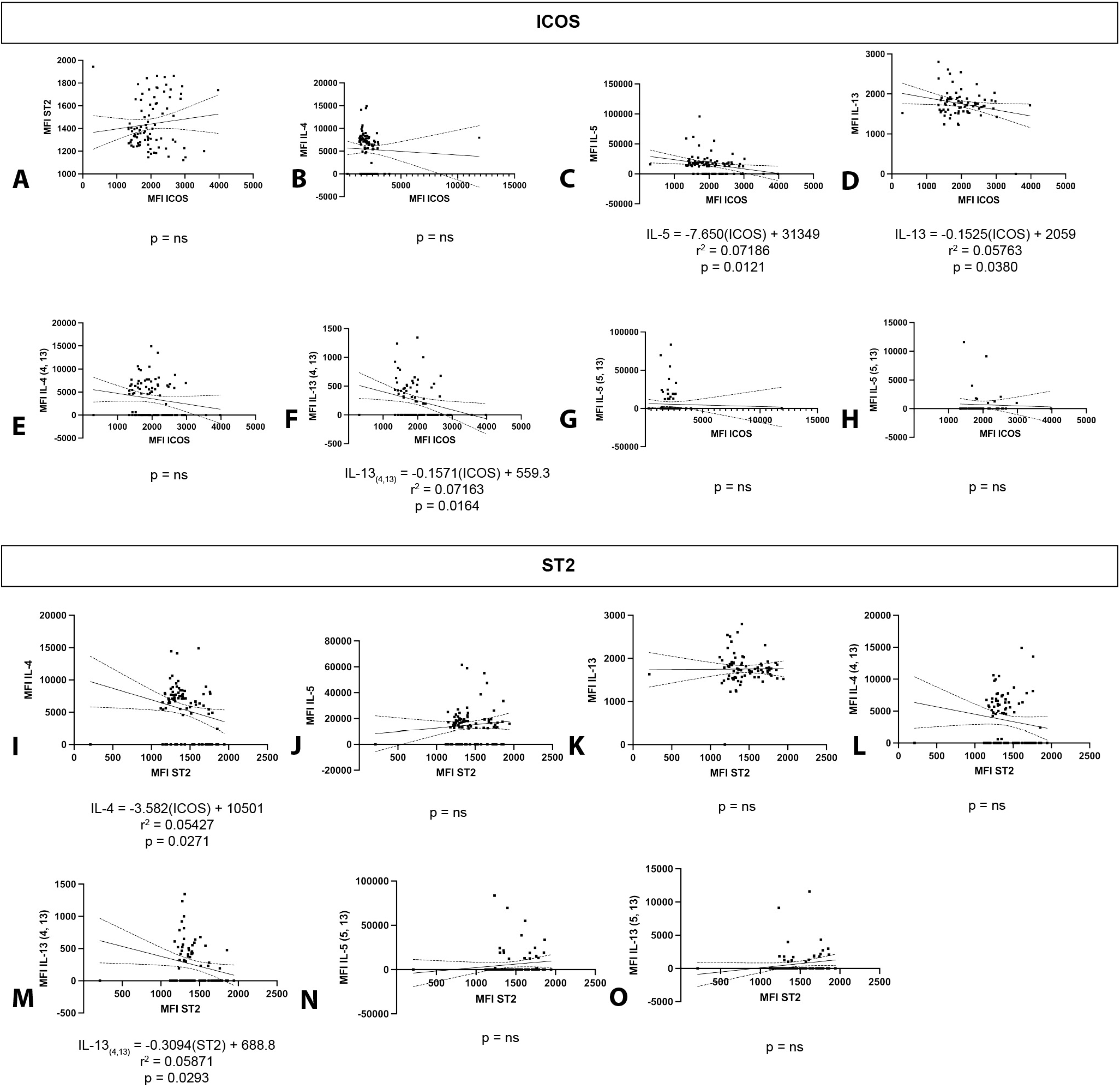
ICOS is an accurate activation marker for brain-resident ILC2s. Lymphocytes of young (6-months) and aged (24-26 months) male and female C57Bl6/J mice were isolated, stimulated overnight with PBS, IL-7, IL-2 and IL-25, IL-33, or PMA, and investigated for type 2 cytokine expression using flow cytometry. Single linear regressions were graphed with 95% confidence intervals for ICOS against ST2 (A), IL-4 (B), and ILC2_4,13_ (C-D) and ST2 against IL-4 (E) and ILC2_4,13_ (F-G). Only significant models were reported with r^2^ and p-values.

### Cell surface markers are correlated to cytokine expression

ILC2s receive and transmit information from many different cell types. Because of this, there is a constant reshuffling of cell surface marker expression and cytokine production to understand their environment; therefore, it is vital to understand how these signals interact and which signals best anticipate ILC2 function.

We used Spearman correlation (Figure 10A) to understand how cell surface markers (ICOS and ST2) and cytokine states (IL-4, IL-5, IL-13, and their various combinations) correlated. Because the inherent factors (tissue, sex, aging) and stimulation were categorical variables, they could not be included in the correlation. Based on the Spearman correlation, the means of most variables positively correlate with each other except for ICOS and IL-4 and IL-5, respectively. To further understand how cell surface markers and cytokine expression influence each other, multiple linear regressions were performed.

**Figure 10.**
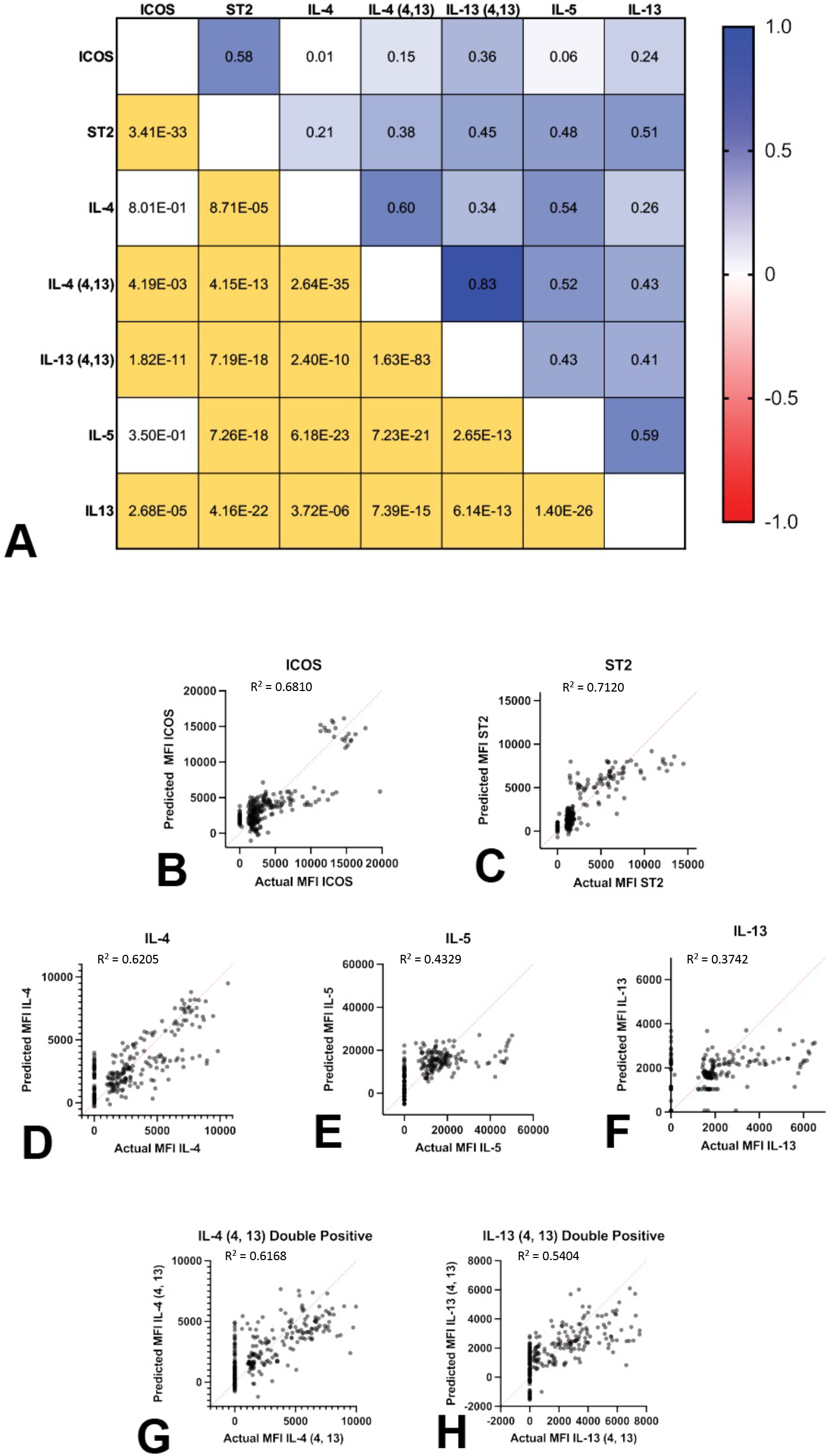
ILC2 expression requires complex interactions to mediate function. Immune cells were isolated from spleen, lung, gut, and brain from young (6-months) and aged (24-26-months) male and female C57Bl6/J mice. Cells were stimulated overnight with PBS (negative control – denoted 1), PMA (positive control – denoted 5) or 20ng/mL IL-7 (denoted 2); IL-2 and IL-25 (denoted 3); or IL-33 (denoted 4). Median fluorescence intensity from stimulated ILC2s were used in a Spearman correlation (A) to understand the correlation between extra- and intra-cellular ILC2 markers. The replicative means are on the right, while p-values are on the left. Significant p-values are denoted in yellow. Multiple linear regressions plotted actual versus predictive outcomes for ICOS (B), ST2 (C), IL-4 (D), IL-5 (E), IL-13 (F), IL-4 of IL-4/IL-13 double positive ILC2s (G), and IL-13 of IL-4/13 double positive ILC2s (H) to include categorical data, such as: stimulation, aging, and tissue type. Complete statistics are reported in Table 2.

**Table 2.**
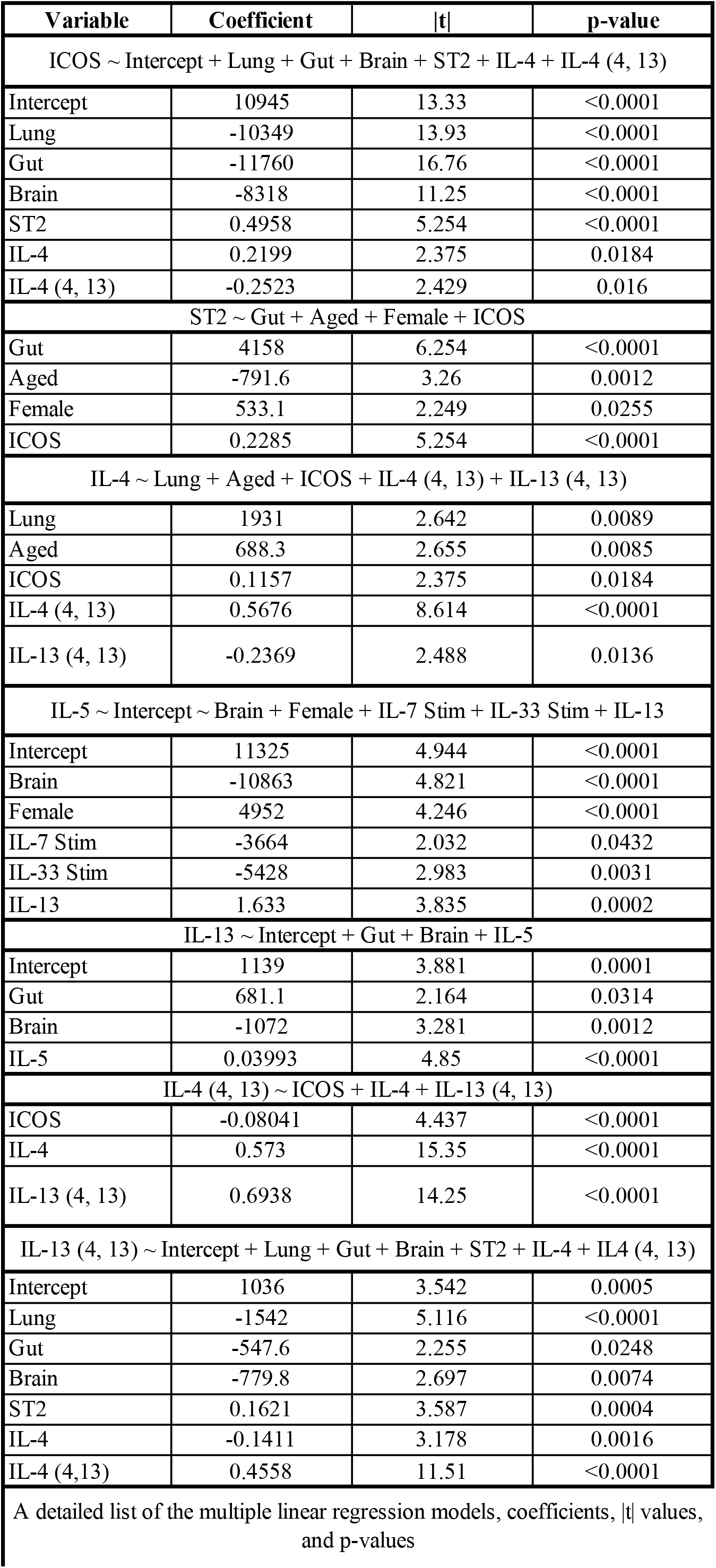
Multiple linear regression model statistics

| Variable | Coefficient | \|t\| | p-value |
| --- | --- | --- | --- |
| ICOS ~ Intercept + Lung + Gut + Brain + ST2 + IL-4 + IL-4 (4, 13) |  |  |  |
| Intercept | 10945 | 13.33 | <0.0001 |
| Lung | -10349 | 13.93 | <0.0001 |
| Gut | -11760 | 16.76 | <0.0001 |
| Brain | -8318 | 11.25 | <0.0001 |
| ST2 | 0.4958 | 5.254 | <0.0001 |
| IL-4 | 0.2199 | 2.375 | 0.0184 |
| IL-4 (4, 13) | -0.2523 | 2.429 | 0.016 |
| ST2 ~ Gut + Aged + Female + ICOS |  |  |  |
| Gut | 4158 | 6.254 | <0.0001 |
| Aged | -791.6 | 3.26 | 0.0012 |
| Female | 533.1 | 2.249 | 0.0255 |
| ICOS | 0.2285 | 5.254 | <0.0001 |
| IL-4 ~ Lung + Aged + ICOS + IL-4 (4, 13) + IL-13 (4, 13) |  |  |  |
| Lung | 1931 | 2.642 | 0.0089 |
| Aged | 688.3 | 2.655 | 0.0085 |
| ICOS | 0.1157 | 2.375 | 0.0184 |
| IL-4 (4, 13) | 0.5676 | 8.614 | <0.0001 |
| IL-13 (4, 13) | -0.2369 | 2.488 | 0.0136 |
| IL-5 ~ Intercept ~ Brain + Female + IL-7 Stim + IL-33 Stim + IL-13 |  |  |  |
| Intercept | 11325 | 4.944 | <0.0001 |
| Brain | -10863 | 4.821 | <0.0001 |
| Female | 4952 | 4.246 | <0.0001 |
| IL-7 Stim | -3664 | 2.032 | 0.0432 |
| IL-33 Stim | -5428 | 2.983 | 0.0031 |
| IL-13 | 1.633 | 3.835 | 0.0002 |
| IL-13 ~ Intercept + Gut + Brain + IL-5 |  |  |  |
| Intercept | 1139 | 3.881 | 0.0001 |
| Gut | 681.1 | 2.164 | 0.0314 |
| Brain | -1072 | 3.281 | 0.0012 |
| IL-5 | 0.03993 | 4.85 | <0.0001 |
| IL-4 (4, 13) ~ ICOS + IL-4 + IL-13 (4, 13) |  |  |  |
| ICOS | -0.08041 | 4.437 | <0.0001 |
| IL-4 | 0.573 | 15.35 | <0.0001 |
| IL-13 (4, 13) | 0.6938 | 14.25 | <0.0001 |
| IL-13 (4, 13) ~ Intercept + Lung + Gut + Brain + ST2 + IL-4 + IL4 (4, 13) |  |  |  |
| Intercept | 1036 | 3.542 | 0.0005 |
| Lung | -1542 | 5.116 | <0.0001 |
| Gut | -547.6 | 2.255 | 0.0248 |
| Brain | -779.8 | 2.697 | 0.0074 |
| ST2 | 0.1621 | 3.587 | 0.0004 |
| IL-4 | -0.1411 | 3.178 | 0.0016 |
| IL-4 (4,13) | 0.4558 | 11.51 | <0.0001 |

A detailed list of the multiple linear regression models, coefficients, |t| values, and p-values

Multiple linear regression was used to test how the inherent factors, cell surface markers, and cytokine expression influence predicted outcomes. For simplicity, all multiple linear regressions will only report the models. Comprehensive statistics are reported at Table 2. The models included: ICOS ∼ Intercept + Lung + Gut + Brain + ST2 + IL-4 + IL-4 (4, 13), R^2^ = 0.6810 (Figure 10B); ST2 ∼ Gut + Aged + Female + ICOS, R^2^ = 0.7120 (Figure 10C); IL-4 ∼ Lung + Aged + ICOS + IL-4 (4, 13) + IL-13 (4, 13), R^2^ = 0.6205 (Figure 10D); IL-5 ∼ Intercept ∼ Brain + Female + IL-7 Stim + IL-33 Stim + IL-13, R^2^ = 0.4329 (Figure 10E); IL-13 ∼ Intercept + Gut + Brain + IL-5, R^2^ = 0.3742 (Figure 10F); IL-4 (4, 13) ∼ ICOS + IL-4 + IL-13 (4, 13), R^2^ = 0.6168 (Figure 10G); and IL-13 (4, 13) ∼ Intercept + Lung + Gut + Brain + ST2 + IL-4 + IL4 (4, 13), R^2^ = 0.5404 (Figure 10H). Based on these models, we infer that certain cell surface markers are not only mediated by the immediate tissue milieu but possibly under communication with other tissues and influenced greater by sex differences or aging. Further studies are required to elucidate mechanisms of ILC-body communication axes.

## Discussion

In this study, we wanted to investigate ILC representation in various tissues and how they are influenced by sex and aging. Previous studies indicated ILCs represent 0.01-0.1% of circulating lymphocytes in humans (23), but here, we show that ILCs comprise 0.2-10% of lymphocytes, mainly rising in aging. Moreover, when comparing percentages between males and females, males have higher ILC representation and more variability between tissues than females. But in both instances, the brain contains the highest representation of ILCs. When considering the brain compared to the other tissues we investigated (e.g., spleen, bone marrow, and lung), the brain is considered immune-privileged and more likely relies on innate immunity like microglia and astrocytes for immunologic censorship (24). The bone marrow is the site of hematopoiesis possibly requiring fewer ILCs and the spleen and lung have high concentrations of T cells (25, 26); therefore, they may not require as much innate censorship/redundancy as other tissues. Moreover, we were able to identify ILC preference based off the variables we investigated. In the young male spleen, ILC1 > ILC3 > ILC2, but in aging ILC1 > ILC2 = ILC3. For the young and aged male spleen and brain: ILC2 > ILC1 = ILC3. The only predisposition for a particular ILC type in females is in the brain, where ILC2 > ILC1 = ILC3 and holds true in aging for all lymphocytes. There has been ample research to indicate that males favor Th1 responses, while females favor Th2 responses because of hormones, where androgens suppress, and estrogens enhance Th2 responses(12, 14, 27, 28). There is still debate on the influence of sex hormones on ILCs, though in places likes the uterus, ILCs are thought to be more sensitive to sex hormones. Moreover, when considering sex, males are ILC1/ILC3 dominant, while females are ILC2 dominant. This is supported by literature that states males favor Th1/Th17 type responses, while females favor Th2 type responses; this effect is closely linked to hormonal sex in young mice (14, 29) but has yet to be elucidated in aging mice.

Further studies need to investigate sex hormones on ILCs and their representation in different tissues. Moreover, because of sex hormone flux in aging, particularly in females, it is imperative to elucidate this phenomenon in aging.

ILCs are plastic depending on the immediate environment of the cells (17). In aging, there is a chronic, low-grade pro-inflammatory response, termed inflammaging, but has not been investigated in ILC representation. Because of the close interdependence of ILC on each other, we wanted to understand the homeostatic balance of ILC representation in aging and sex differences. We determined that tissue type and sex influence ILC representation, but aging was not the strongest influencer of ILC modulation. This implies that ILCs do not require numeric compensation against inflammaging but modulate their function instead. This is partially supported by our data, but further research is required to understand the complete function of all ILCs.

With ILC plasticity being a crucial ILC characteristic, we wanted to address if the cells displayed plasticity in their markers. We used a stringent lineage gate to filter out as many known, uniquely defined cells from our query, including T and B cells, myeloid cells, antigen presenting cells, granulocytes, endothelial cells, and epithelial cells. We then used NK1.1 for ILC1s, ST2 for ILC2s, IL-23R for ILC3s, and NKp46 for cytotoxicity (18). Reducing dimensionality with UMAPs and quantifying marker-specific MFI, we concluded that there was little plasticity indicated in our assay except for the aged brain, which also yielded the highest ST2 MFI value. In general, the markers consistently clustered with themselves on the UMAPs, though the brain tends to favor four main cluster groups that are influenced by sex and aging and not strongly effected by stimulation. When quantifying the markers, splenic ILCs indicate the most plasticity with NK1.1 expressed on all three ILC types in variation though the highest expression is on ILC1s; moreover, this effect is maintained in aging. We recognized a similar phenomenon in the brain, particularly where ILC1s are the main cell type to express ST2, which suggests a non-committed, plastic ILC1/2 phenotype. The bone marrow and lung maintained their ILC lineage commitment. The role of the spleen in mediating and maturing T and B cells to antigen presenting cells may necessitate flexibility in ILC populations to influence immune cell polarization that is not readily seen in sites like the lung and brain. The brain’s immune privilege may require plasticity in ILCs for fast responses to brain injuries or maintaining brain health. Future studies would need to investigate cell signaling, localization, and ILC preference in each tissue.

When compared to the other tissues, ST2 expression is highest in the brain. In mice, IL-33 is constitutively produced and in the highest concentrations in the central nervous system (30, 31), which may explain the high concentration of ST2 there. In our study, we discovered that ST2 is highly expressed on brain ILC1 in young females that then decreases in aging where ST2 expression is equal between ILC1s and ILC2s and males and females. The increase of ST2 on young, female ILC2s may be to negate the low numbers of ILC2s in the brain, which is not required in aging, but more experiments are needed to understand this phenomenon. Moreover, brain ILCs are the most cytotoxic compared to other tissues, which may be mediated by the immune privilege in the brain and lack of robust immune responses by sentinel microglia and patrolling T cells in the cerebrospinal fluid. However, the spleen and bone marrow have sex differences in ST2 expression, where males express more ST2 than females. Future studies should probe both hormonal and chromosomal sex differences on systemic ILCs using systems like the four-core genotype to understand why ST2 expression is different in males compared to females, particularly in the spleen and bone marrow which aren’t classically known as a hormone-producing organs.

As ILC2s are the unique, anti-inflammatory ILC type, we wanted to explore their cytokine potential thoroughly. After stimulating ILCs overnight with various conditions, we used flow cytometry to quantify IL-4, IL-5, and IL-13 and elucidate how these cytokines are differentially expressed in tissues, sex, and aging. We determined that there are ILC2 subtypes based on cytokine expression. We are not the first to propose ILC2 subtypes, as Fung et al., 2020 (20) also found ILC2 subtypes based on RNA-seq data, but they only investigated aged females. ILC2s favor single cytokine expression, but there are small subsets that can concurrently express IL-4, IL-5, and IL-13 in various combinations. No matter the cytokine expression profile, there were cytokine production differences in each tissue as anticipated. The spleen and lung were equally able to express all cytokine types and double cytokine types. Of the tissues studied, the brain was the least likely to have double or triple positive cytokine expressing ILC2s and favored IL-4 expression over the other cytokines. The lung was the only tissue with triple positive ILC2s. ILC2_4,13_ were represented in all tissues of this study and better represented in aged brain; however, ILC2_4,5_ were more common in the spleen and lung whether young or aged. Even though IL-4, IL-5, and IL-13 are regulated by GATA3, there are other transcriptional mechanisms that require exploration to fully understand ILC2 cytokine predispositions, especially if they are to be used translationally. Mechanistic studies could elucidate pharmaceutical targets for ILC and possibly be used in their adaptive counterparts to understand immunosenescence.

To further probe ILC2 complexity, we wanted to elucidate predictors of extra- or intra-cellular markers in a systems biology approach. Except for ICOS with IL-4 or IL-5, all other factors correlated with each other in our Spearman correlation. Using multiple linear regression, we were able to determine how each factor influenced the other expression values. As anticipated, ICOS and ST2 are not influenced by the stimulation variables that we used. This would suggest that ILC2s do not change their cell surface markers at the overnight timepoint perhaps from immune exhaustion or communication with the other cells they were incubated with. Of all the cytokines investigated, only IL-5 was influenced by different stimulation types. Furthermore, the models we generated were able to fit the data between 35-80%, which continues to suggest a multi-variate and complex approach to ILC2 expression and activation. As mentioned before, future studies will need more unbiased investigations into the transcriptome and proteome of ILC2s in tissue, sex, and aging to fully understand how ILC2s can be used therapeutically.

To take a more simplistic approach, we did simple linear regressions to determine if the extracellular markers ICOS and ST2 could be used as ILC2 activation markers. We showed that, when significant, ICOS was negatively correlated to its target in the spleen and gut. This was opposite in the lung and brain where ICOS was positively correlated with its targets. ST2 yielded positive correlations in the spleen, lung, and brain, but negatively correlated in the gut. Of the organs investigated, the gut requires a different marker to predict ILC2 output. This also holds true for IL-5 expression as none of the markers investigated correlated with its expression. ILC2s present many distinct cell surface markers like KLRG1, IL7Ra, CD117, and TSLPR that require further investigation as predictive activation markers on ILC2s. Future studies should also investigate the if this translates to human ILCs to be used clinically for different diseases. This could be utilized over human cytokine ELISAs that are potentially more time consuming and costly than cell surface staining, especially in such a rare cell population and even more so in the blood.

In summary, this study investigated how tissues, sex, and aging influence ILC representation. Moreover, we probed cytokine profiles and predictive activation measures for ILC2 potential. Future studies would benefit from more unbiased approaches to understand transcriptome and proteome changes in ILCs to enhance ILC foundational information. Similar studies would need to be repeated, particularly in human blood, to understand ILC basic concepts before manipulation in different disease states.

